# MazeMaster: an open-source Python-based software package for controlling virtual reality experiments

**DOI:** 10.1101/2020.01.27.921148

**Authors:** Alexander Bexter, Björn M. Kampa

## Abstract

In the last 15 years, virtual realities have revolutionized behavior experiments in particular for rodents. In combination with treadmills, running wheels, or air-floating balls, the implementation of a virtual reality (VR) provides not only the opportunity to simultaneously explore behavior and neuronal activity in head-fixed animals under nearly natural conditions, but also allows full control over the visual sensory input presented to the animal. Furthermore, VRs can be combined with other sensory modalities such as auditory, tactile or olfactory stimuli. Despite the power of using VRs in animal experiments, available software packages are very limited, expensive and lack the required flexibility to design appropriate behavior and neurophysiology experiments. For this reason, we have developed the versatile, adaptable and easy to use VR environment *MazeMaster*, an open-source, Python-based software package for controlling virtual reality setups and behavior experiments. The software package includes a graphical user interface (GUI) and can be integrated into standard electrophysiology and imaging setups even by non-programmers. Ready-made behavioral experiments such as multisensory discrimination in T-mazes are already implemented including full control for reward supply and bias correction. For more individual setup designs, the modularity of *MazeMaster* allows more programming-affine users to extend the software with potentially missing features. With *MazeMaster*, we offer a free and easy-to-use VR controller that will facilitate the implementation of VR setups in scientific laboratories. In addition, *MazeMaster* allows the design and control of common head-fixed rodent behavior paradigms with extensive acquisition of meta-data required for reproducible VR experiments. The *MazeMaster* VR package, therefore, offers a collaboration tool for reproducible research within and across neuroscience laboratories according to the FAIR principles.

## 1 Introduction

Performing behavior experiments while simultaneously recording brain activity in rodents can be difficult if the animal is free to move or has to change its position constantly. An attempt to solve this problem is to head-fixate the animal and place it in a virtual reality. This has originally been realized in rats ([1]) and later also in mice ([2]; [3]) and gerbils ([4]). Following the most common procedure, the virtual reality is rendered on a computer and then presented to the test subject on either standard LED monitors (e.g. [5]; [6];) or with the help of a projector on a dome-like screen (e.g. [1]; [7]). It is crucial that the setup screens cover a large area of the visual field, which itself is relatively big due to the lateral arrangement of the eyes in mice and rats. Movements within virtual space are realized with a treadmill, running wheel or an air-suspended floating ball on which the animal can run. A wheel is commonly used in linear virtual tracks, in which moving in one dimension is sufficient, whereas the floating ball is used for two-dimensional movements, mostly in complex mazes or open spaces. An advantage of a virtual maze is the perfect control over the sensory input to the test subjects. Visual, auditory and tactile stimuli can be presented at high temporal and spatial precision ([8]; [9]; [10]). A study by [11] studied whisker-guided locomotion by using movable walls on both sides of the animal to simulate tunnel walls. Similarly, [10] used varying sandpapers on rotating cylinders to mimic different wall textures. Providing auditory stimuli in a virtual environment together with visual cues to test navigational behavior was realized by [12] or [9]. Recently, a similar approach has also been used to create a virtual maze for olfactory cues ([13]).

Together with the locomotion of the animal, movements of the pupils and whiskers can be tracked ([10]). It is also possible to modify the environment while the animal is moving in it. Furthermore, it is not limited in space, providing the possibility to use test fields of infinite size. Rodents can navigate in such virtual spaces like a straight tunnel (e.g. [1]; [3]; [5]; [6]) or more complex mazes like a T-maze (e.g. [14]; [15]) or even stacked U-mazes ([4]). A lot of recent reports have shown that the recorded cortical activity differs dramatically between periods of quiescence or rest and locomotion ([16]; [17]; [18]). For example, the visually evoked firing rate of most neurons in the primary visual cortex of awake mice was more than doubled when the mouse transitioned from standing still to running ([19]). Place- and grid-field responses in the hippocampus and medial entorhinal cortex were found as well when using virtual environments ([2]; [7]; [5]; [6]). Therefore, it might be reasonable to allow the animal to run during the behavior experiment. One obvious way to implement this would be to place the behavior experiment in a virtual reality.

Different VR implementing software solutions were created and used during the last years. Existing game engines could be applied to create VR environments. For example, Harvey et al., 2009 used the software *Quark* (http://quark.planetquake.gamespy.com) to build a virtual linear track, which uses the open-source Quake2 game engine (id software). The Blender Game Engine (www.blender.org) is another popular software tool to create three-dimensional environments. Combined with the Blender Python API, it was driving the virtual reality system in a study by [20]. [4] used Blender for modeling three dimensional mazes and exporting them afterward. The control of the VR was done with another software (Vizard Virtual Reality Toolkit; WorldViz, http://www.worldviz.com) and additional custom-made software for communicating the infrared mouse data. Further studies used the 3D graphics engine Ogre (http://www.ogre3d.org) and OpenAL [21] or OpenSceneGraph (FlyVR, http://flyvr.org/) [22]. However, those approaches often lack a user-friendly interface to create and control virtual environments and have to be combined with additional scripts to read the input. Another extensive VR software package (and similar to *MazeMaster*) is ViRMEn (http://pni.princeton.edu/node/4071) which was created and published by Dmitriy Aronov ([23]). It is a MATLAB-based software, which also consists of a graphical user interface (GUI) in combination with a (OpenGL) 3D graphics engine. Yet, none of those packages offer the possibility to create and run a whole experiment, including the control of all behavioral parameters and the application of multisensory virtual cues. There are also commercial solutions available that provide the hardware and software to run several experiments in virtual reality. Those systems often use TFT monitors or spherical dome screens for visual stimulation and balls, wheels, or belts as running devices. Limitations of those systems are often the restricted experiment types, therefore those systems can be used for recreating existing experiments with established, standardized environments. However, since the software is not open-source, the experimental design cannot be changed and individualized. Similar restrictions might apply to the hardware, which often cannot be exchanged freely.

Importantly, these packages lack the possibility of embedding the VR environments into a fully-functional behavioral experiment and providing the user with the complete tools to plan, create, run, and save such an experiment. To overcome those limitations of existing software, we developed the *MazeMaster* software package. *MazeMaster* is designed as an all-in-one software package, which allows the user to create new experiments, including VR environments, in a few minutes. The combination of a graphical drawing-board-like maze building interface (*MazeBuilder*) and an interface for automated creation and monitoring of complete behavioral and neurophysiological experiments is an outstanding feature of this toolbox. It is possible to extend the functionality to present stimuli of various modalities and even combine these to create a multisensory virtual environment. As an example, a demo is provided in which air puffs are implemented in a virtual corridor to combine tactile and visual cues. Custom scripts can be added and started at individual time points during an experiment to flexibly add any additional functionality to the program. The software package is an approach to standardize similar experiments by using the same open-source tools ([24]) as also recommended by the FAIR principles ([25]). Since a lot of experiments in the field of behavioral neuroscience use similar experimental setups and designs, it would be favorable to share common software toolboxes. A lot of resources can be saved by preventing long development processes to create own customized software, but instead, build on already existing and approved frameworks covering the required functionality. The provided software toolbox (*MazeBuilder*) is free to use and open-source. Researchers are invited to use the toolbox under the open-source license and further extend its functionality. The whole software is written in Python, which is also free of charge and commonly used. Figure 1 shows an overview of a VR setup used with MazeMaster (panel A and B) as well as an example view from within a virtual maze (panel C).

**Figure 1:**
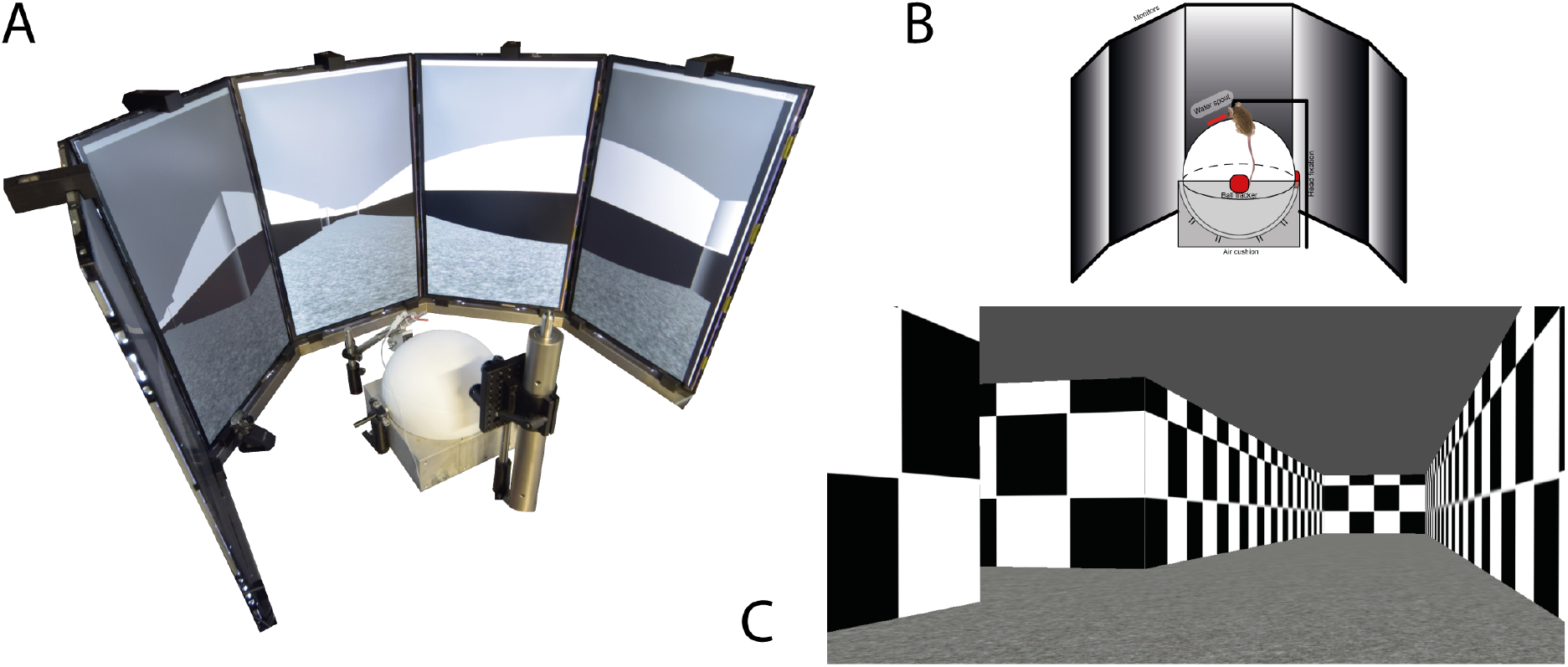
A) Setup of a virtual reality with five monitors in a semicircle with a ball as input device. B) Schematic drawing of the setup in A. C) View from within a virtual Maze.

## 2 Software Description

*MazeMaster* comes as a bundle that connects a 3D engine (Blender game engine, included in the package) with a powerful maze experiment design and control tool. With this, it is possible to create and modulate complex mazes, simple tunnels or other forms of three-dimensional constructs. The tool can be used to design and conduct behavioral experiments for mice and rats but also human participants. *MazeMaster* consists of several modules, which are combined within a graphical user interface (GUI). This GUI is provided to help the user with creating and monitoring virtual environments and experiments. There is no coding knowledge required to use it. The toolbox already includes parts for setting up a 3D environment, run an experiment and save the corresponding data. No further software is necessary. To combine the VR behavior experiments with electrophysiological recordings or imaging of brain activity digital triggers can be introduced for the communication of *MazeMaster* with the acquisition software. Usually, experiments are performed in closed-loop with the animal starting and finishing each trial. In this case, *MazeMaster* would act as a master controlling any further data acquisition as a slave. However, experiments can also be performed under external control for open-loop configurations.

### 2.1 Environment

The software is written in Python 3.7. Blender files are created with Blender v2.79. So far, the software is only running on a Windows system. The MazeMaster toolbox can be installed and used as a pre-compiled file on a Windows PC. No further installation of Python or additional modules is required.

#### 2.1.1 Python Modules used in the creation progress

Several existing python modules were used to create MazeMaster. The GUI was created with the *tkinter* module. Network communication was realized via the asynchronous socket service handler *asyncore*. For hardware control, the module *nidaqmx* was used, which is the interface for all National Instruments hardware. To use it, the NI-DAQmx drivers from National Instruments have to be installed. All modules are listed in supplemental table S2.

### 2.2 Hardware

The software is compatible with different kinds of hardware. A complete list of hardware parts that were used with the software so far can be found in the supplemental table S1. Theoretically, any device can be used for displaying the maze (e.g. computer screens, projectors), since the window can be scaled to any size. Nevertheless, the angle of displaying the maze has to be changed according to the relative position of the subject. So far, there are pre-configured settings for one or two monitors, as well as an arrangement in a semicircle (c.f. [6]). The latter includes monitors, but also projectors that project the image onto a dome. We tested a setup with five monitors in a semicircle, using a *ASUS R9 280 series* graphics board.

To acquire the animal locomotion data several configurations have been implemented for a linear or spherical treadmill as input devices measuring the movement of the animal. As a linear treadmill, we have used a custom build styrofoam wheel, which uses an incremental encoder (Kübler) to measure the rotation of the wheel. As a spherical treadmill, we used a 20 cm diameter styrofoam ball floating on an air cushion allowing the head-fixed animal to run on it and move in 2D. The rotation of the ball was measured with two optical computer mice (Logitech Anywhere MX). Both configurations can be selected in the MazeMaster GUI. Additional input devices can be added, by extending the tracking module. For the rotation encoder as an input device and also for the digital triggers a National Instruments (NI-6323) card with a breakout box (BNC-2110 box and SCB-68A box) was used. Other devices like an Arduino could be implemented in the future to replace the NI components. We have implemented specific digital triggers for water valves to supply a water reward in the behavioral experiments. Furthermore, we have introduced an interface for a lick detection as required in behavior tasks. It is possible to trigger other devices such as imaging or electrophysiology devices via a set of generic digital triggers, whose timings can be configured separately. A typical setup is shown in figure 2 B. The connectivity between the different parts is shown by the black lines and arrows.

**Figure 2:**
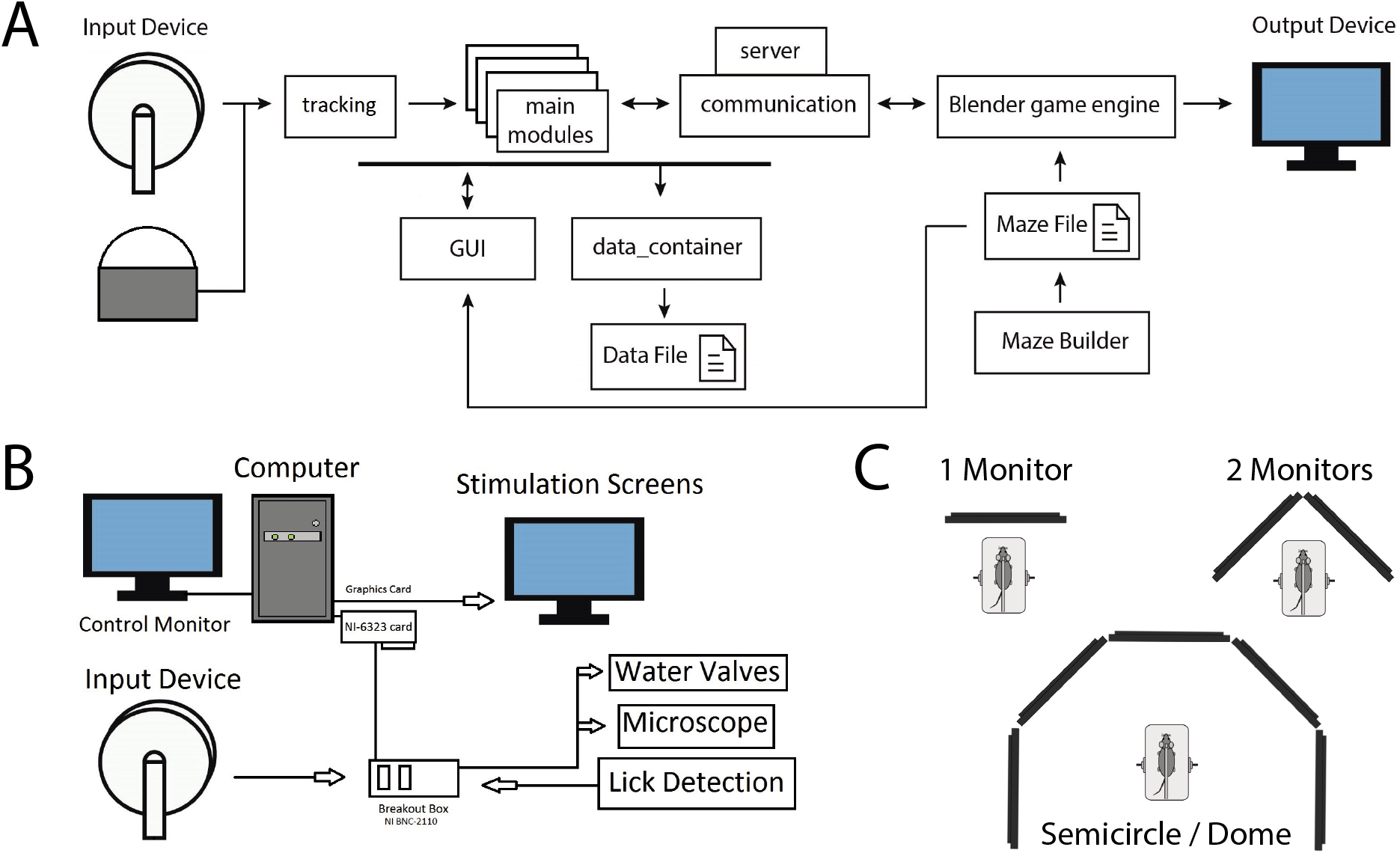
**A)** Sketch for general stream of data. Arrows indicate the direction of data flow and communication. Main modules represent all other modules, which are not mentioned separately. The communication module uses the server module to establish a connection and communicate with the blender game engine. **B)** Sketch for general hardware parts of a setup to use with *MazeMaster*. Arrows indicate the direction of data flow and communication. Water valves, microscope, and lick detection are used in most setups but can be freely extended with further devices. **C)** Possible modes for displaying the virtual reality on three different arrangements of monitors or screens. The display mode can be selected in the settings.

### 2.3 Software Architecture

The program is divided into several modules, containing classes with different functions. An overview of the most important modules and the stream of data is shown in figure 2 A. Since *MazeMaster* is designed to be extendable, more modules with new classes can be created to add new functions to the program.

The main part which connects all included modules is the *MazeMaster* GUI. With this main GUI everything can be controlled, including the *MazeBuilder* and the Blender interface. All *MazeMaster* Python modules are imported into the main script from where the initialization of all class objects is done and the GUI is started. The already implemented modules often use an individual GUI window to add new functionality. Those functions are active, as long as the corresponding GUI window is open. Those windows are connected to the *MazeMaster* window top menu through which they are opened.

New mazes can be created by using the *MazeBuilder*, which comes with a drawing board-like GUI. This allows the creation of an extensive variety of mazes that can be saved and archived as *.maze* files. The *MazeBuilder* is a standalone program and can also be used without the GUI of *MazeMaster*.

The input from the connected devices is processed by the tracking module and transmitted to the main modules. A new device can be created by adding a new class to the module, so far there are classes for a ball and a wheel. All information is relayed to the blender game engine via the communication class. For this, a socket connection is used with the help of the server class. All data is stored in a data container and saved at the end of every trial. Most information is also displayed in the *MazeMaster* GUI. The maze file, which is created with the help of the *MazeBuilder*, is read by the maze-building script (*createmm.py*, not shown) and used by the game engine.

#### 2.3.1 Integration with Blender Game Engine

For creating and displaying the maze on the monitors, the Blender 3D game engine (www.blender.org) is used. It is provided together with *MazeMaster* as a standalone version. A blender file (*virtualMaze.blend*) is run in the *blenderplayer*, a standalone executable from blender to run the Blender game engine. A python script (*createmm.py*) is run when the connection to Blender is established with the help of the server module. This script creates the maze from the maze file as a three dimensional model in Blender and configures all parameters in Blender for the environment. After that, it saves all changes to the Blender file (*virtualMaze.blend*). For all further communication, the network connection, which was started by *server.py*, is used. This connection uses an asynchronous socket for communication. Similar to using multi-threading, the asynchronous socket allows *MazeMaster* to continue running while communicating in the background without having to wait for finishing the network tasks.

### 2.4 Functionality

#### 2.4.1 MazeMaster Main Graphical user interface

With the GUI, it is easy to set up and configure an experiment. It is divided into different sections (see Figure 3). The top menu *Windows* opens several additional windows, which are not open by default (for a list see supplemental table S3). The drop-down menu *View* lets the user configure and save the default positions of the different windows. In the *General Settings* section the user can define different sets of settings for the experiment control, as well as the file saving directory (see section 3.3). Those sets of settings are saved as experiments and associated tasks, which can be freely created and loaded by the user. The *Server Control* section allows the user to start the Blender Game Engine and connect *MazeMaster* to it. The *Server Status* indicator shows the status of the connection to the graphical engine (online, waiting for connection, offline). The *Disconnect* button stops the connection to the engine and closes the session. The *Cues* section lists all loaded visual cues and lets the user load new cues and delete not used ones. Those cues are loaded and created from image files, which should be square-shaped. Different sets of visual cues can be loaded, too. The chosen cues can be presented at defined locations within the virtual reality indicated by the cue identifier number. The cues are either constantly visible (*Cues on Walls*) or invisible until a cue trigger is present or until the subject reaches a specified location in the maze where the cues are then flashed on the maze walls (*Flash Cues*). If a certain sequence of cues should be presented, there is the option of loading a sequence containing csv file via the *load Sequence* button. This file contains the IDs of cues (those cues have to be loaded with the corresponding IDs in the GUI), which are automatically presented sequentially. The *Trial Control* section allows the user to set up all trial-related settings and provides information on the current experimental status. In this part, most of the relevant behavioral settings can be made. Position tracking can be activated with a certain frequency (only needed without a wheel as an input device). A set of up to four digital triggers can be configured with different time points. The trial starting mode can be either set to directly (automatic start) or triggered (running start). A water reward is set up with a certain valve opening time and probability. The *Add Comment* function further allows adding notes and comments during the experiment similar to an electronic lab notebook (ELN). These notes will be saved together with the experiment data and can be used for automated data analysis routines. Session settings can be set here, too. The user can define the number of trials, the inter-trial-interval, maximal trial duration, and maximal trial length. An entire behavior experiment can be designed and controlled with *MazeMaster* and some common features such as the bias correction, which allows correcting for a side bias in a 2-alternative-choice/ T-maze task, can be set. Also, an air-puff can be triggered manually and the air puff time can be set. Movement in the maze can be done automatically with a given speed (constant) or a loaded profile (variable) from a recent trial (from trial data file) by enabling autorun. Also the length of one square in the maze can be set here, changing the speed of the input device. The *Maze Info* panel lists information regarding the currently loaded maze.

**Figure 3:**
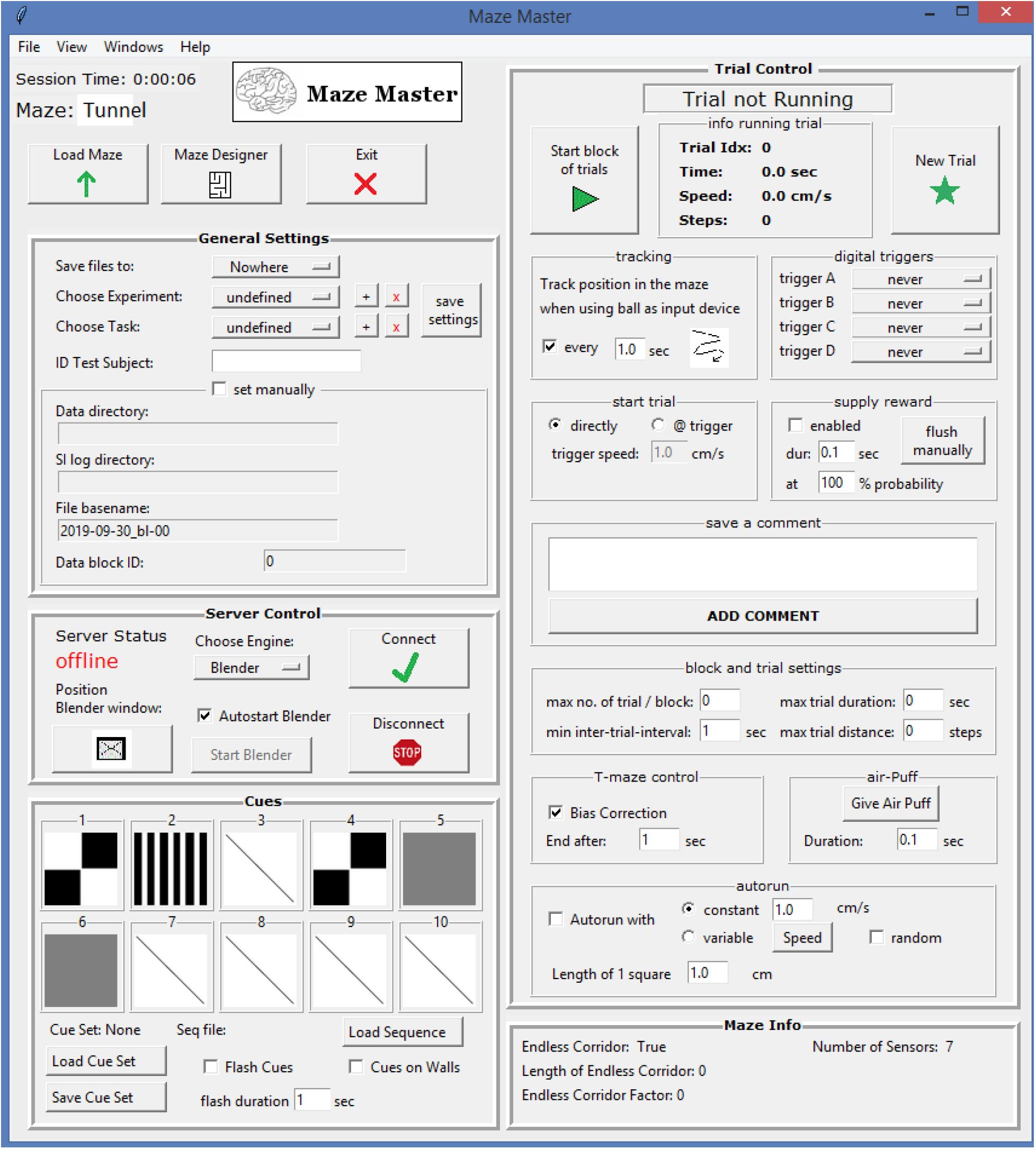
The main graphical user interface (GUI) for *MazeMaster*. It is divided into several panels, which allow the user to set up and configure different kinds of experiments in a virtual maze.

#### 2.4.2 The MazeBuilder Tool

The mazes used in *MazeMaster* can be of any shape, all kinds of more complex mazes can be created with the *MazeBuilder*. It has to consist of one starting point and at least one endpoint or reward zone. With the *MazeBuilder* tool it is possible to create new mazes by using a drawing board interface. Every element of the maze can be placed and shaped, which includes walls, start and end positions, reward points, teleportation points, and moving walls. It is also possible to create a maze automatically, which is the easiest way to create corridor mazes. This can be done by defining the size of the maze manually by either directly entering the length or by choosing the number of cues or images to show on the corridor walls. In the latter case, the maze is created automatically, matching the number of images. The maze is then saved in the maze folder and can be loaded from *MazeMaster*. It is also possible to change the maze either by opening it again in *MazeBuilder* or opening the file and changing the numbers manually.

##### Starting points

At this point, each trial is started. The virtual user is oriented towards the southern direction of the maze by default.

##### Reward points

When reaching one of the reward points, a reward can be given. The trial ends and the next trial is started after a set inter-trial-interval. The reward amount (usually in milliliter) can be configured and can be provided with a certain reward probability, which is set in the GUI.

##### Teleporter points

This point is the position where the virtual user is moved to when the teleportation feature is used. Teleportation is normally done, after reaching a certain position sensor or at the end of a trial back to the start point.

##### Position Sensors

Position sensors can be placed anywhere in the mazes using the *MazeBuilder*. They are invisible walls and function like light barrier sensors. When the virtual user passes these sensors certain events will be triggered. Those events are:

- Showing visual cues on the maze walls
- Teleport the virtual user
- Place/remove an additional wall in the maze

Figure 4 A shows a virtual tunnel that is used as an endless corridor. The last sensor is used to teleport the user back to the beginning of the tunnel, to create the impression of endlessness. The other cues are used to flash images (cues) on the walls of the tunnel. They are automatically created when using the create (endless) maze by pictures function in the maze builder. For a further description see chapter *events*.

**Figure 4:**
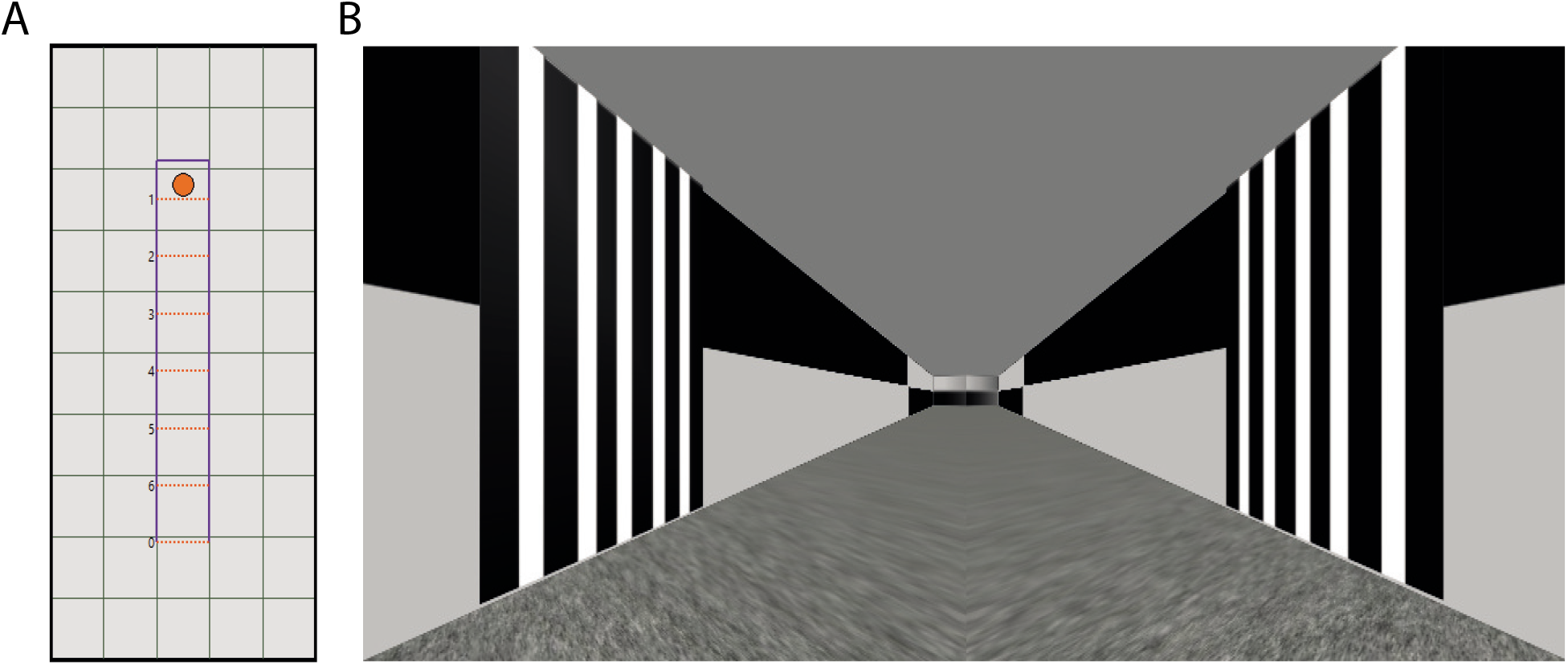
**A)** An example of a virtual tunnel, containing seven position sensors. In this example the last sensor (0) is used for teleportation to the beginning, whereas the other six (1-6) are used for triggering cue presentation. **B)** Example screenshot for a virtual tunnel maze, in which a cue is flashed to the maze walls.

#### 2.4.3 Maze Design: Special maze types

##### T-Maze

A special type of complex maze is the T-Maze. It consists of two arms in which the reward points are placed. Only one of them is shown during the trial, chosen by random. Alternatively, both of them can be made invisible and additional cues can be given to indicate the correct arm of the maze to run into.

##### Corridor Maze

This maze type consists of a linear tunnel. It is functionally similar to complex mazes when a reward point is placed at the end of the corridor. Cues can be used as background texture on the walls. An often used type of corridor maze is an **endless corridor**. In this type of maze, the corridor seems to be endless, the virtual user can move without reaching an end. Nevertheless, there can be a separation into single trials, which are saved and restarted after a certain distance or time. This becomes handy when triggering other hardware like a camera or a microscope, or just to separate the data into repeating parts.

#### 2.4.4 Input Devices

In most cases, the movement of the animal is controlled by either a ball or a wheel. *MazeMaster* offers interfaces for incremental rotary encoders (1-D movement) or movement read by optical/laser computer mice (2-D movement). Which device to use is specified in the configuration window. The connection to the rotary encoder can be established by using a National Instruments card and the appropriate drivers (nidaqmx module for Python). The position of the wheel can then be read as a number, which is depending on the resolution of the encoder (e.g. 360 or 1024 ticks per cycle). The ball can be read by positioning two computer mice on two orthogonal sides. The movement of the ball is then recorded by those optical mouses and the movement is transferred to *MazeMaster*. New kinds of devices can be set up by adding classes to the *input.py* module, inherited from the *Tracking* base class.

##### Movement Modes

Normally, the virtual subject moves accordingly to the input of the encoder (Wheel) or the computer mice (Ball). The length of one square in the maze can be defined in the trial control section. In addition to the normal running, there is the possibility of using the *autorun* mode (control section). This mode can be used either with a constant automatic movement (constant mode), or a movement profile of a recent trial (variable mode). For the latter option, the trial data file has to be loaded to extract the movement data.

##### Starting the trial

A trial can be started either directly or with a running trigger. When the trigger is used, the start of the trial is delayed until a certain speed is reached on the wheel/ball. This mode enables the test subject to start the trial by themselves trough running.

#### 2.4.5 Digital Trigger

To connect and communicate with all kinds of other devices (or software), there are up to four digital triggers to use. They can be individually triggered either once at the beginning of the session, at the beginning of each trial or the end of each trial. The triggers can be used with a NI-card.

#### 2.4.6 Behavioral Features

##### Lick Detection

It is possible to use up to two input signals, which are normally used as licking detection devices. Those two lines are just digital input pulses and therefore can be used with all kinds of devices.

##### Rewards

When a reward area is reached, a water reward can be delivered. This can be done automatically with a certain probability and duration/amount of water. Up to two valves are controlled by this via the NI-card. The signal lines of the valves have to be connected to the ports of the NI board to trigger the opening.

##### Behavior Analysis

When doing a behavior task, it is important to monitor the outcome of all trials, to check for parameters like performance or to detect bias. This can be done in the behavior panel and fits experimental designs based on a 2-AFC task. Experiments are normally done in a T-Maze or the tunnel maze.

##### Air-puffs

An air-puff can be given by clicking the button in the Trial Control section. The valve (digital port) will open for the set duration.

#### 2.4.7 Events

All kinds of events can be triggered automatically when reaching a light-barrier-like sensor in the maze. The rules for programming the sensors are defined in the rules window. Any number of events can be set for each sensor.

##### Visual Cues

The placing of visual cues on the maze walls is done automatically by passing one of the position sensors. Up to 10 cues can be loaded from texture files. The presentation time can be set, as well as the side (left or right wall) of stimulation. The cue appears on the wall at a 45 angle from the point of view on the chosen side.

##### Teleport

Teleportation can be done when passing one of the position sensors. The virtual subject is placed immediately at the teleportation point.

##### Additional walls

Predefined additional walls can be placed in the maze when passing a certain position sensor. Alternatively, predefined additional walls can be removed.

#### 2.4.8 Custom Scripts

User-written scripts can be loaded to add additional functionality to *MazeMaster*. Those scripts are normally written in python or loaded as a batch file and are executed at adjustable time points during an experiment (start of answer period, end of answer period, start of trial, on reward, on error, on missed). With the help of those scripts, optional functions can be added without any limitation.

### 2.5 Blender game engine as the virtual environment

Blender game engine is a component of the modeling software Blender, used for creating real-time interactive content in a realistic 3D environment. The *MazeMaster* package contains a blender file (*.blend*), which is executed with the *blenderplayer* and runs the scripts. The pre-modeled maze from the maze builder is loaded by a custom-written script from within the Blender game engine and is then automatically created as a 3D model. The engine provides realistic physical behavior for the models, which is especially important for collision with physical boundaries in the virtual world, like the maze walls. With the help of this continuous collision detection, the user is stopped by walls, but not by invisible position sensors or reward areas. Theoretically, even movement in height and therefore in the third dimension is easily possible, but so far not used since movements are either one-dimensional along a corridor or two-dimensional within a flat maze. Every maze is built by creating its walls and position sensors at the defined positions. All other elements, like reward areas and visual or sensory cues, are only placed at the defined positions. This means, that those elements can easily be changed (e.g. size, shape, colors, transparency, etc.) in the blender file.

### 2.6 Experimental Data

The data from each experiment session is saved as one general metadata file containing information about the session, one metadata file containing all information about the trials and one data file for each trial. Each occurring event is saved in the data file together with a maze position and a time point. When an encoder (wheel) is used, each movement is tracked and saved. When computer mice are used as an input device, the tracking of position and saving of this data is done at a sampling frequency, specified in the *MazeMaster* GUI. Figure 5 A shows an excerpt from an example *MazeMaster* data file for one trial. The movement of the wheel is tracked in an event-based manner. Each change of the wheel position (steps) is saved as a line in the file, together with the time and virtual coordinates (xpos, ypos) within the virtual environment. All other events are also tracked event-based as shown in the three last lines in the example. First, a reward zone was entered (reward_ID 0), followed by a lick at the left water spout (*lick* column). After that, the left water valve opened for 0.05 sec (*reward* column). Other saved events include passing a position sensor, entering a reward zone, giving a water reward, showing a visual cue and detecting a lick.

**Figure 5:**
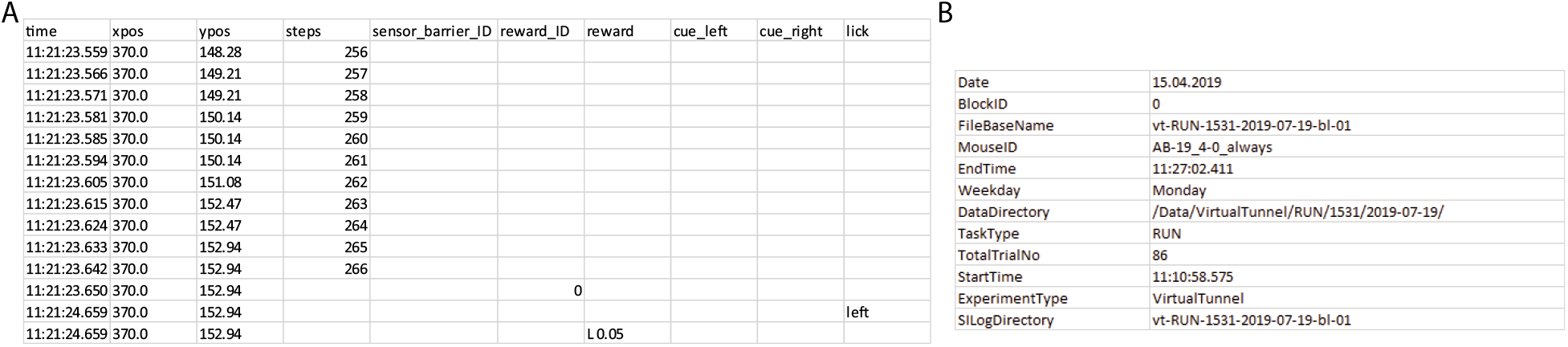
**A)** An excerpt from a *MazeMaster* Data output file. Data rows are collected event-based. Each row has a timestamp with the corresponding position in the virtual environment (*xpos, ypos*). If a wheel with an encoder is used, each step is recorded as a row with the current step counter in the column *steps*. All other events are listed in the rest of the columns. In this example, the reward area with the ID *0* has been reached, with a lick on the left lick detection, recorded in the following line. After that lick, a reward was given also on the left side with a valve opening time of 0.05 seconds. **B)** A metadata file, containing all information about a session.

All information regarding the trials can be found in the metadata file for the trials. There, the trial duration, target-stimulus and response location, the replay file path are saved and if comments have been entered by the user during the trial. The comment itself can be found in the metadata file for the session as an electronic log or notebook (ELN). The data is saved as a csv file. Other formats would also be possible. The common file format can be easily opened in spreadsheet processing programs like Excel but also in Python or Matlab. Data is stored in an experiment-, task- and subject-specific file path. This path is created automatically when the task is selected and a name for the test subject entered. The folder structure is hierarchical, ordered first by the experiment, then the task, test subject and finally the date of recording. The filename is a combination of the task, together with the date and the block ID. This number indicates the number of recordings (blocks) on the same day and is incremented automatically. This method should help to maintain a structure for saving and organizing data files. The collected metadata further allows reproducible experiments and automated data analysis routines which are an important requirement for the FAIR principles ([25]; [26]). The file path can be changed manually, too. The metadata file contains information about the experiment. An exemplary data set is shown in figure 5 B.

## 3 Setting up an experiment

The following section describes, how to set up a new experiment, starting from building a new maze to set up stimuli and trial settings.

### 3.1 Create a New Maze

With the *MazeBuilder* tool it is possible to create new mazes by using a drawing board like interface. Every element of the maze can be placed and shaped, which includes walls, start and end positions, reward points, teleporting points and moving walls. It is also possible to create a maze automatically, which is the easiest way to create corridor mazes. This can be done by defining the size of the maze manually by either defining the length directly or by the number of pictures to show on the corridor walls. In the latter case, the maze is created automatically, matching the number of images. The maze is then saved in the maze folder and can be loaded from *MazeMaster*. It is also possible to change the maze either by opening it again in *MazeBuilder* or by hand, opening the file and changing the numbers manually.

#### Placing a wall

A new wall is created by clicking on an empty node on the drawing board, this will create the starting point of the wall. To set the wall you have to click on another node as the endpoint. The wall is created and you can continue with the next one, while still in creating mode. To end the creating mode click the right mouse button. All walls are marked in purple.

#### Deleting a wall

You can delete a wall by simply clicking on an existing wall with the right mouse button.

#### Placing reward points

Reward points are positioned by clicking on them and then click on the position where it should be placed. The number of reward points can be set by clicking on the minus and plus buttons.

#### Placing the start point

The start point is positioned by clicking on the button and then click on the position where it should be placed. The virtual player will face the southern direction of the maze when the trial starts.

#### Placing the teleporter point

The teleporter point is positioned by clicking on the button and then click on the position where it should be placed. After a teleportation event is triggered, the virtual player will be transported instantly to this teleporter point.

#### Placing sensors

A position sensor is placed by first clicking on the button and then on a starting point, followed by an endpoint of the sensor wall.

#### Placing false walls

A false wall is placed by first clicking on the add sensor button and checking the *False Wall* checkbutton. The wall is placed like a sensor by defining a starting point, followed by an endpoint of the wall.

#### Creating an endless maze

This maze type consists of a linear tunnel. It is functionally similar to complex mazes when a reward point is placed at the end of the corridor. Cues can be used as background texture on the walls. An often used type of corridor maze is an endless corridor. In this type of maze, the corridor seems to be endless, the virtual user can move without reaching an end. Nevertheless, there can be a separation into single trials, which are saved and restarted after a certain distance or time. This becomes handy when triggering other hardware like a camera or a microscope, or just to separate the data into repeating parts. The normal way of creating an endless corridor is to set the number of pictures, which should be on the wall. This determines its length and can even be used without using any pictures in the end. The endless maze factor sets the factor with which the length will be multiplied to give the impression of an endless maze (default = 6). After entering those values a click on the button *create endless maze* will create the corridor. **Textures** Textures for the walls can be loaded by using the buttons in the *Textures* section. The *Background* button loads a texture for the maze walls. Textures for ceiling and floor can be loaded by clicking one of the corresponding buttons. The textures (image files) should be square in size.

#### Saving the maze

After finishing the design, a name should be entered. After that, clicking the save button will save the maze in the maze folder.

### 3.2 Load a maze

A maze can be loaded by clicking the Load Maze button. It will be opened and displayed in the maze window. The name of the maze is shown above the load maze button on the top.

### 3.3 General Settings

The place where to store the data can be defined here. There are some presets from which to choose, which can be changed in the settings (*Windows* - *Settings*). There are presets of settings that can be defined and saved for each task separately. First, the experiment has to be chosen, to which the task is assigned. You can add new experiments by clicking the plus button next to the dropdown menu. The same can be done with tasks, which are then assigned to the currently chosen experiment. Tasks and experiments can be deleted by clicking the red x next to the dropdown menu. Be aware that if you delete an experiment, all assigned tasks are deleted as well. The currently selected settings are saved to the chosen task by clicking the save settings button. For identification of the dataset, an ID can be entered for the test subject. The filename and directory are then created automatically out of the experiment, task, and ID. Alternatively, it can be set manually by checking *set manually*.

### 3.4 Trial Control Settings

In this section, all the settings regarding the trial can be set.

#### Start tracking

If this setting is enabled, the path the virtual subject took in the maze is tracked by a line with the entered frequency.

#### Digital trigger

There are four digital triggers for any device (e.g. camera, microscope), which are either triggered directly at the start of each trial or only once at the beginning of the first trial.

#### Start trial

With this setting, you can switch between an automatic start of each trial and a movement triggered start. When the triggered start is chosen, the start of each trial is delayed until the given speed is reached for at least one second.

#### Supply reward

If this setting is enabled, a water droplet is given, when the virtual subject enters a reward Area. The amount can be set via the duration of the valve opening. In addition, a probability can be set, with which the reward will be given each trial. The *flush manually* button will open/close all valves.

#### Block and trial settings

In this block, the maximal number of trials can be set. After reaching this number, the program will automatically stop running. The maximal trial duration determines the amount of time after the trial is terminated automatically. The same goes for the maximal trial distance, which determines the number of steps after the trial is terminated automatically. A zero means infinite in all cases, the trial/experiment will not stop automatically. The entry field minimal inter-trial-interval sets the delay time between trials.

#### T-maze control

The T-maze, or in general all 2-alternative-forced-choice experiments in *MazeMaster*, can use a simple bias correction, to correct for a side bias. You can enable this by clicking the checkbutton. Furthermore, the time can be entered in *End after*, after which the trial ends when entering one of the arms in a T-maze.

#### Air-puffs

An air-puff can be given by clicking the button in the Trial Control section. The valve (digital port) will open for the set duration.

#### Autorun

Normally, the virtual subject moves accordingly to the input of the encoder (Wheel) or the computer mice (Ball). The length of one square in the maze can be defined here. In addition, there is the possibility of using the autorun mode. This mode can be used either with a constant movement (constant mode), or a movement profile of a recent trial (replay or variable mode). For the latter option, the trial data file has to be loaded to extract the movement data from by clicking the *Speed* button. For the randomization of those recent running profiles, simply check the *random* checkbutton.

### 3.5 Loading Cues

The cues are shown in the cues section on the left side. Ten cues can be loaded in total to show on the maze walls. A new cue can be loaded by clicking on the image/placeholder of the cue. An already loaded cue is replaced with the new one. You can delete cues by right-clicking on the cue image. All loaded cues can be saved as a set of cues, which can be loaded afterward. For this, use the *Load Cue Set* and *Save Cue Set* buttons. Cues can be shown in two different modes: Flashed or as the texture on the wall. When the flashed mode is used, the images will be flashed on the walls at the sensor’s positions for a certain duration, which can be set here as well. The sensor’s positions are normally automatically created from the *MazeBuilder* so that the images are distributed throughout the length of the corridor. The cues are flashed from left to right/ascending order, which means, that the cue number zero will always be the first one. The number of cues should match the number of sensors, but ignoring the sensor number zero (teleportation sensor). That means there should be one more sensor than cues.

When the *cues on Walls* mode is chosen, the cues will be as a texture on the walls, lined up from left to right. Again, the number of cues should match the length of the corridor. Normally one cue equals one square (or sensor) on the maze window.

#### Varying sequences of cues

It is possible to change the cues from trial to trial automatically. A predefined list of cues can be loaded via the Load Sequence button. This list should be a .csv file with one column named “cue”. In this column, all the cues should be listed with their index numbers according to the cue indices in *MazeMaster*. An often-used method is to load more cues than sensors and automatically change the cues, which should be shown, via the sequence file. Each row should contain one cue index number, the program will simply assign the next cue in the list to the next sensor and after finishing the trial, it will continue with the next position in the list, assigning it to the next sensor again. You can find an example sequence file for alternating cue triplets in the doc folder.

### 3.6 Further Settings

#### Change Input Device

Go to *Windows* - *Input devices* and select *wheel* or *ball* and configure those devices.

#### Adding Scripts

Open the script window by clicking *Windows* - *Scripts*. New scripts can be added by clicking the *add script* button. After loading a python (.py) or batch (.bat) file, trigger have to be defined for executing those scripts.

#### Set Maze Settings

Open the settings window by clicking *Windows* - *Maze Settings*. Here, the scaling of the maze can be defined, as well as certain rules for restarting the trial (after reaching the reward or after teleportation). Additionally, sensors can be configured as reusable.

#### General Settings

The window *Windows* - *Settings* let the user control the display mode of the VR, the blender window position, as well as the data directory and all digital ports (for triggers, reward, lick detection, etc.)

#### Behavior Control

To use the behavior module, simply open the window *Windows* - *Behavior*

### 3.7 Starting the Experiment

To start the experiment, the Blender game engine has to be started and connected to *MazeMaster*. This can be done by first clicking on the connect button in the *Server Control* section. When the *Autostart Blender* option is marked, Blender will start automatically when clicking the connect button. Otherwise, Blender can be started by clicking the *Start Blender* button. Blender will open a window and connect automatically to *MazeMaster*. When the connection is successfully set up, the server status will change to *connected*. The first trial can be started by clicking on the *Start block of trials* button in the *Trial Control* section.

## 4 Example Experiments

The following section shows four example experiments in virtual environments, which all have been tested thoroughly with mice. Each experiment can be implemented by following the steps in the documentation file (doc.pdf) in the folder *examples*. This folder also contains all other files, which are necessary to run the experiment. The created data in the example experiments has been briefly analyzed and is shown in the following sections.

The first experiment is a navigation experiment in a virtual tunnel. The second experiment is a navigation experiment in a virtual maze. The third experiment is an endless tunnel, where cues are flashed on the walls, as the mouse is running through it (Data not shown, but the example can be found in the “Virtual Endless Tunnel” folder). The last experiment shows a whole behavioral task in a virtual tunnel, where visual stimuli where shown on the tunnel walls and were discriminated by a mouse.

### 4.1 Navigation in a Virtual Tunnel

This simple tunnel experiment is best suited as a first experiment, to train mice to run in a virtual environment. The virtual reality setup with a ball as the input device (see Figure 6 A) was used. Mice had to run from one end of the tunnel, where the start position was placed, to the other end of the tunnel, marked by a reward area. The position in the virtual space was tracked with a frequency of one Hz. The position data was extracted from a *MazeMaster* data file for each trial. An example of such a file can be found in figure 5 A, where positions are saved with the corresponding time points.

**Figure 6:**
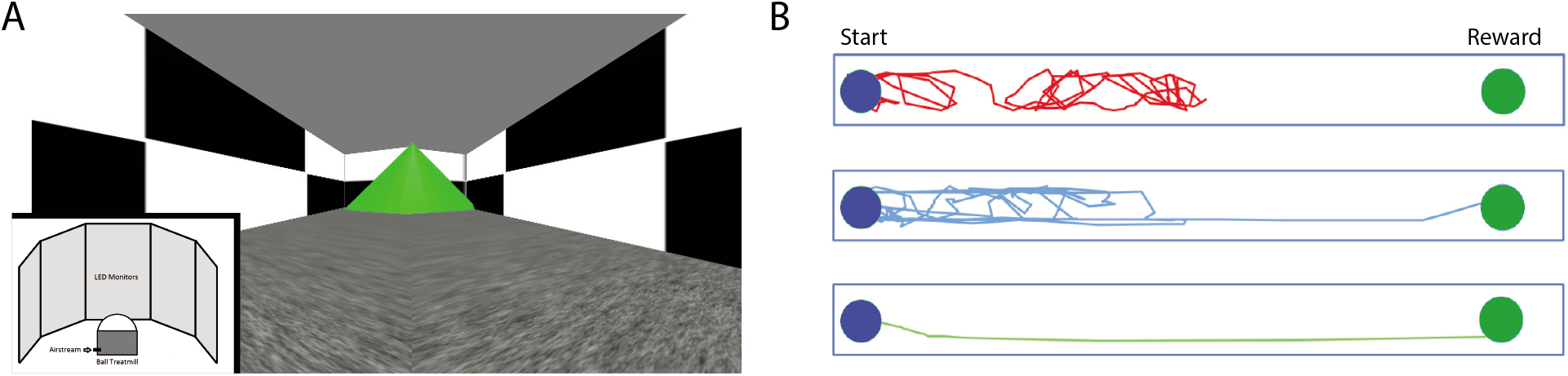
A) View from within the virtual tunnel with the reward area at the end. Inlet shows a sketch from the setup, which uses a ball as the input device. B) Tracked movement in the tunnel during three example sessions from different time points during the experiment. The green circle depicts the starting point, the blue circle the reward area.

The change in running behavior indicates, that mice were able to learn the movement in a virtual environment and optimize their running behavior over time. The recorded running positions in the tunnel can be easily visualized, too. Figure 6 C shows the path the mouse took in the visual tunnel in three example trials from three different sessions.

This example experiment can be found as a demo for *MazeMaster* in the folder “Virtual Tunnel”. The documentation file contains a step-by-step description of how to create and run the whole experiment.

### 4.2 Navigation in a Virtual Square Maze

Mice did a navigation experiment in a virtual reality setup with a ball as the input device, to allow for movement in two dimensions through the maze. A reward area was placed in a virtual square-shaped maze. The mouse had to find this area, starting from the same starting point in each trial. There were different possible routes to the target area, which were of different lengths. Mice should learn to find the target area, using the shortest route over time. The route the animal took was tracked, as well as the time it took to reach the target area. The different possible routes were separated for the analysis. Mice learned successfully to run to the reward area, using different routes, which are pictured in figure 7 A. The example figures were created using the position tracking with a frequency of one Hz. Positions are saved with the corresponding time points in the data file (see figure 5 A).

**Figure 7:**
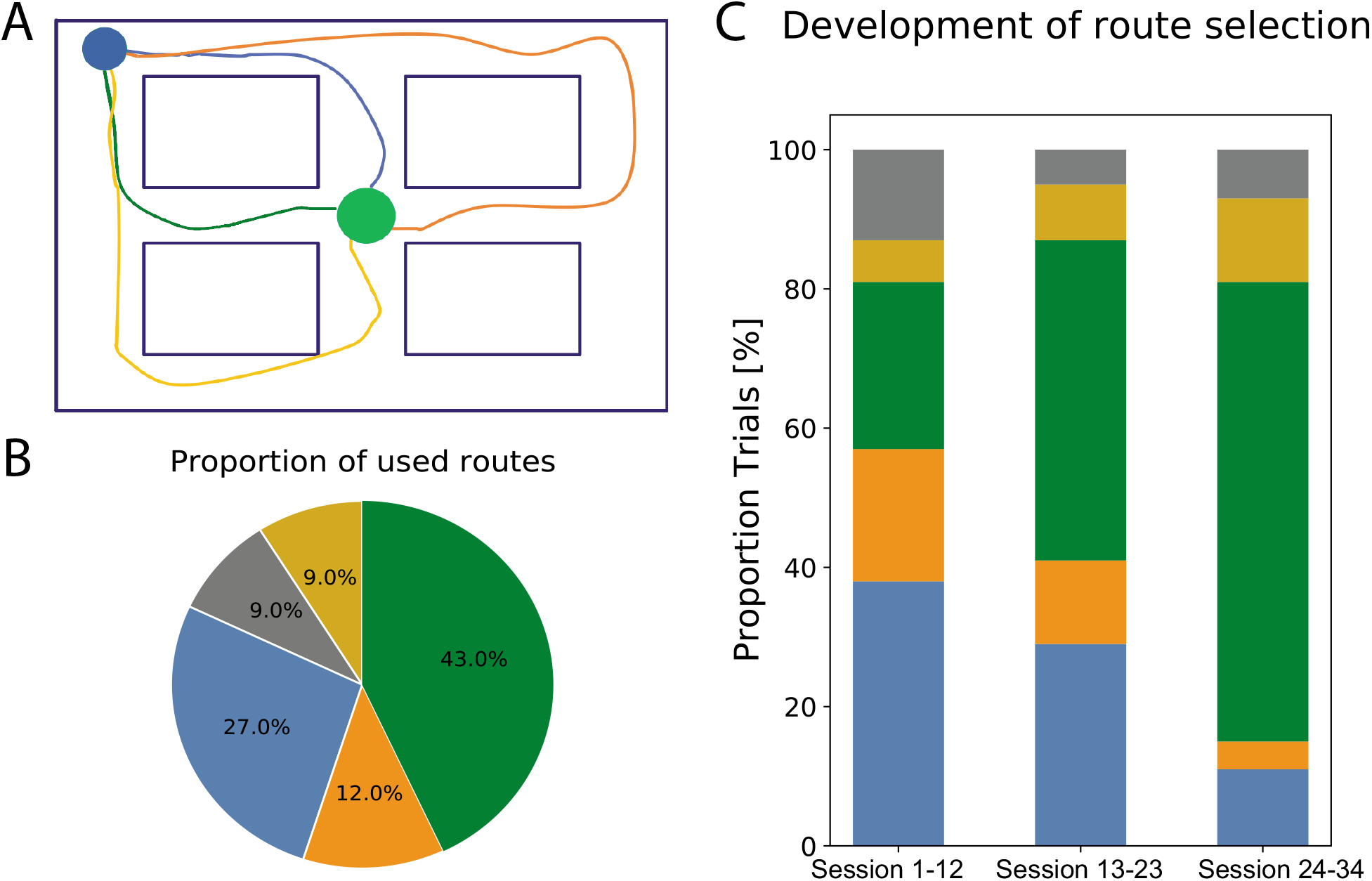
Movement behavior in a complex square maze. A) Different paths are shown, which the mouse took to get to the reward area in an example session. The blue circle shows the starting position, the green circle represents the reward zone. The colors of the routes match the colors in B) and C). B) The proportion of routes the animal took in total. The corresponding routes are shown in A). The grey part represents other routes, which are not shown with a color. C) The development of the proportion of routes the animal took over the course of the experiment. The total of 34 meetings were divided into three blocks. Grey part represents other routes, which are not shown with a color.

This example experiment can be found as a demo for *MazeMaster* in the folder “Virtual Square Maze”. The documentation file contains a step-by-step description of how to run create and run the whole experiment.

### 4.3 Endless Tunnel with Flashing Images

In this experiment, a mouse was running on a wheel in an endless corridor (see figure 6 A for the setup). Images were appearing on the walls (flashing), as it moved through the tunnel (see figure 4 B). The tunnel appears to be endless since the animal is seamlessly teleported back to the beginning when the last position sensor is reached. The length of the tunnel is adapted to four wall images in a row without any gap in between. This set of images is provided and can be loaded as a set of cues. Each cue presentation is triggered by a position sensor, thus together with the sensor for teleporting the animal back to the beginning, there are five sensors in total. This example experiment can be found as a demo for *MazeMaster* in the folder “Virtual Endless Tunnel”. The documentation file contains a step-by-step description of how to run create and run the whole experiment.

### 4.4 Multisensory Stimulus Discrimination Task in a Virtual Tunnel

A virtual tunnel was used in a behavioral experiment, where two sets of visual cues were projected onto the walls. The mouse was running on a wheel until the end of the corridor to the reward zone. The head-fixed mouse had to discriminate between two sides of the tunnel and detect the side with more cues (white bars) by licking at one of two water spouts, associated with the two sides/tunnel walls (see figure 6 A for the setup). The task resembles the task introduced by [15]. In this example, simple bars where projected onto the walls as cues. But it is possible to change the models of the cues in Blender so that they are of any three-dimensional shape like for example towers (as in [15]). Furthermore, air puffs or acoustic tones from two sides can be added in addition to the visual cues for multisensory stimulation. To run this mode in a virtual corridor, the *flash tunnel* module is used. *MazeMaster* was used in this 2-alternative forced-choice experiment, to create the experiment, monitor the realization and save the data. An excerpt from the behavioral data is shown in figure 8 A. The trial data file contains trial information like start- and stop time, the target side, the answer side and the ID of the stimuli presented in the trial. It also saves the name and file path of a replay file (in *autorun* mode), if one was used for the trial. The list provides an overview of trial-specific data and metadata, like if there was a comment created in a specific trial.

**Figure 8:**
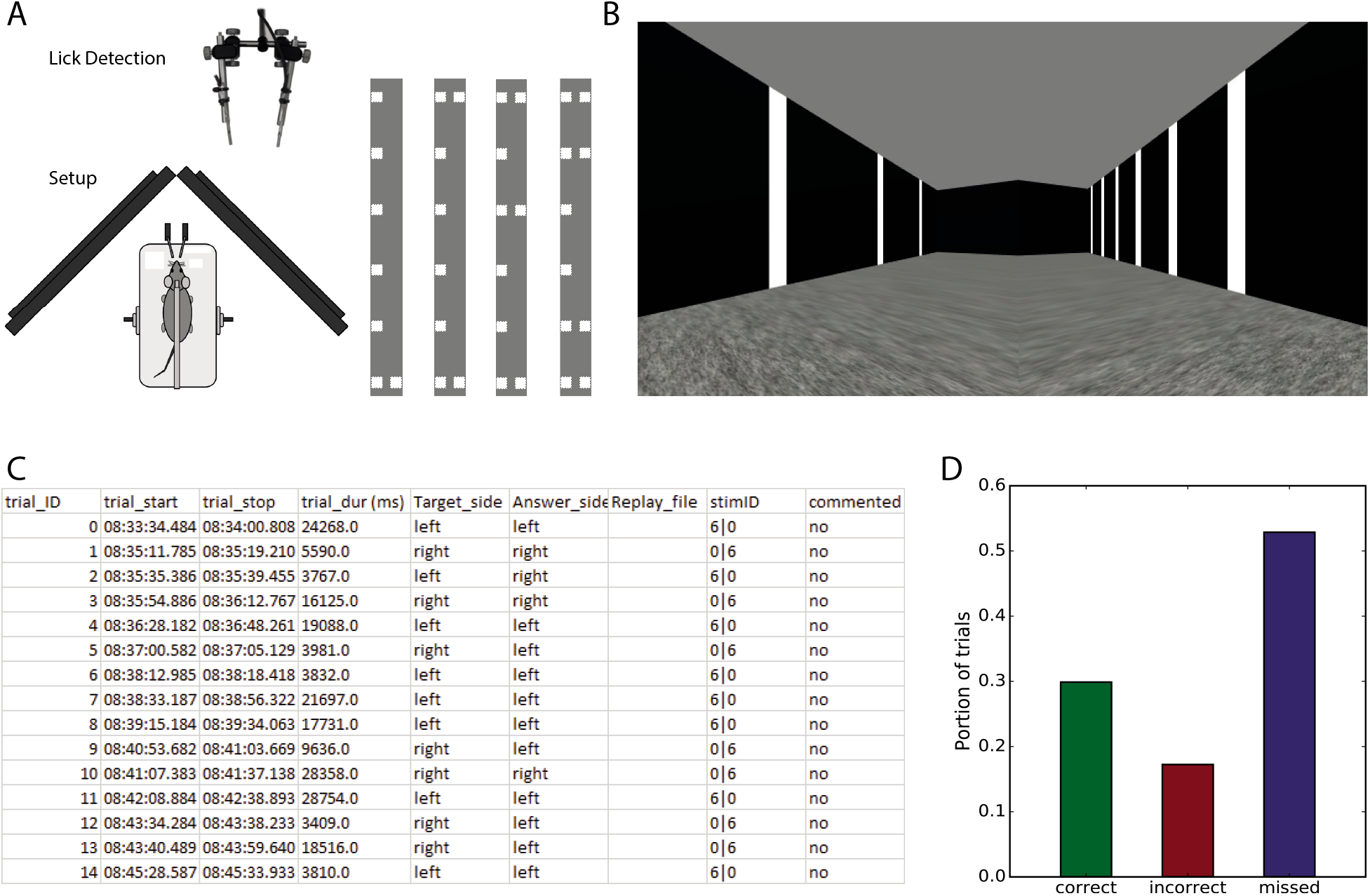
A) Left: Schematic drawings of the Setup using a wheel as the input device. It includes two lick spouts for answering right or left. Right: Structure of the virtual tunnel with different sets of cues on the walls (four different distractor sets). B) View from within the virtual tunnel in an example trial with a distractor of three white bars against the target of six white bars. C) Excerpt from a MazeMaster trial overview file, including trial parameters and behavioral responses. D) Distribution of correct, incorrect and missed trials during an example session, using data from a trial overview file.

The data was used to extract the information and generate multiple plots out of that. Figure 8 B shows the performance of an animal during the session in A. Additionally, data from the lick detection and running data recorded by a linear decoder in a running wheel was recorded.

This example experiment can be found as a demo for *MazeMaster* in the folder “Virtual Tunnel Discrimination Task”. The documentation file contains a step-by-step description of how to run create and run the whole experiment.

## 5 Discussion

The software package provided here delivers many options for performing experiments in a virtual maze. It can be used by programmers and non-programmers alike and offers user-friendly tools for creating test environments and conducting experiments in those virtual spaces. It has numerous possible options for connecting hardware devices and is therefore suitable for many different experimental approaches. In addition, in cases in where the software is not able to fulfill the requirements for a certain kind of experiment, *MazeMaster* is extendable in a modular way. Another, easier option is to use scripts, which can be executed automatically at predefined time points. Those files are mostly python scripts, which are simply run during the experiment and which can contain all kind of additional code. At the moment the software is only running on Windows systems. It would make sense to extend this functionality to Linux based operating systems in the future. We chose Blender as an engine for three-dimensional virtual mazes because it is nicely accessible with the help of the Blender Python application programming interface (API). It is also freely accessible and free of charge. *MazeMaster* was written in Python and is therefore freely accessible. This can be an advantage compared to software packages, that are based on commercial software (e.g. MATLAB).

A disadvantage when using the three-dimensional rendered virtual maze is, that there is a delay of up to 30 ms between the input signal (e.g. of the encoder of the wheel) and the reaction in the virtual reality and update of the monitors. For most behavior and imaging experiments however it might not be a problem. In timing-critical measurements synchronizing triggers can be recorded additionally for post-hoc synchronization of the recorded data.

The connection with the special hardware (input devices, digital trigger/input) should be extended to other devices in the future. For this, the input device class can be extended in a modular way. Additionally, to control the valves for water supply and air puffs or use digital signals as output or input a National Instruments board is used. This could be extended to use devices like an Arduino, a Raspberry Pi or similar single-board controllers.

Together *MazeMaster* offers a versatile open-source toolbox for conducting behavior experiments in virtual reality. The implemented features allow a broad range of experiment and maze designs and provide a direct link for simultaneous imaging and electrophysiological recordings. Further customization is easy to incorporate by the user and for a common benefit of the community.

## 6 Current Code Version

**Table 1:** current code version.

|  |  |
| --- | --- |
| Code version | 1.0.0 |
| Permanent link to code repository | https://github.com/BRAINLab-Aachen/MazeMaster |
| Programming Language | Python 3.7 |
| Legal Code License | GNU GPLv3 |

## Author Contributions

AB and BMK have designed the software package and experiments. AB has written and tested the software. AB has performed and analyzed the experiments. AB and BMK have discussed the results and wrote the manuscript.

## Acknowledgments

The authors want to thank Lyuba Zehl and Mara Caltapanides for their help with the design and writing of the software and Gerion Nabbefeld and Magdalena Robacha for test runs and helpful feedback.

