## Supplemental for "MazeMaster: an open-source Python-based software package for controlling virtual reality experiments"

### Table S1. List of materials

**Computer setup**

2 Fujitsu B24T-7 Monitors for VR

2 NI-6323 cards, National Instruments

1 NI BNC-2110 box, National Instruments

1 NI SCB-68A box, National Instruments

1 Graphic board for VR, ASUS R9 280 series

**Lick-detection and water reward**

1 Mount for the plastic tubes, Thorlabs parts

2 Plastic tubes

2 Silicone tube, Roth

Flexible tubes for water reward, Braun

2 Piezoelectric ceramic bi-morph elements

1 Professional moving magnet preamp, Dynavox (TC-750)

2 two port solenoid valve for water, SMC (VDW22LA)

**Polystyrene wheel**

1 polystyrene wheel, 20 cm diameter, Rayher

1 incremental encoder, Kübler

1 Mount for the wheel, Thorlabs parts

### Table S2: List of modules

- sys
- asyncore
- time
- tkinter
- numpy
- csv
- os
- datetime
- configparser
- socket
- rawinputreader
- ast

- win32api
- PIL
- shutil
- random
- tarfile

### Table S3: Additional windows in Maze Master

- Tunnel Flashes
- Behavior
- Input Devices
- Maze Settings
- Rules
- Scripts
- Settings
